# Reliability of resting-state EEG modulation by continuous and intermittent theta burst stimulation of the primary motor cortex: A sham-controlled study

**DOI:** 10.1101/2023.05.12.540024

**Authors:** Andrei Rodionov, Recep A. Ozdemir, Christopher S.Y. Benwell, Peter J. Fried, Pierre Boucher, Davide Momi, Jessica M. Ross, Emiliano Santarnecchi, Alvaro Pascual-Leone, Mouhsin M. Shafi

## Abstract

Theta burst stimulation (TBS) is a form of repetitive transcranial magnetic stimulation designed to induce changes of cortical excitability that outlast the period of TBS application. In this study, we explored the effects of continuous TBS (cTBS) and intermittent TBS (iTBS) versus sham TBS stimulation, applied to the primary motor cortex, on modulation of resting state electroencephalography (rsEEG) power. We first conducted hypothesis-driven region-of-interest (ROI) analyses examining changes in alpha (8-12 Hz) and beta (13-21 Hz) bands over the left and right motor cortex. Additionally, we performed data-driven whole-brain analyses across a wide range of frequencies (1-50 Hz) and all electrodes. Finally, we assessed the reliability of TBS effects across two sessions approximately 1 month apart. None of the protocols produced significant group-level effects in the ROI. Whole-brain analysis revealed that cTBS significantly enhanced relative power between 19-43 Hz over multiple sites in both hemispheres. However, these results were not reliable across visits. There were no significant differences between EEG modulation by active and sham TBS protocols. Between-visit reliability of TBS-induced neuromodulatory effects was generally low-to-moderate. We discuss confounding factors and potential approaches for improving the reliability of TBS-induced rsEEG modulation.

## Introduction

Theta burst stimulation (TBS) is a form of repetitive transcranial magnetic stimulation (rTMS) designed to produce long-lasting modulation of neural excitability^1^. Inspired by animal studies of synaptic plasticity^2^, conventional TBS consists of 3-pulse 50 Hz bursts of TMS administered every 200 ms. In the intermittent TBS (iTBS) protocol, 2 s stimulation trains are spaced by 8 s inter-train-intervals (ITIs), with 600 pulses in total typically being administered, resulting in increased excitability as indicated by increases in TMS motor-evoked potentials (MEPs) for up to 30 min post-stimulation. In the continuous TBS (cTBS) protocol, the patterned rTMS is applied continuously over 40 s, resulting in decreases in MEP amplitudes for up to 60 minutes ^3^. This bidirectional pattern-specific modulation of neural excitability by TBS is thought to be linked to synaptic plasticity in the human cortex ^4^ via induction of long-term potentiation and long-term depression mechanisms. In comparison to conventional fixed-frequency rTMS, TBS possesses several advantages, including shorter stimulation times and lower stimulation intensities ^5^.

Although initial studies ^1,6^ of TBS showed expected neuromodulatory effects, more recent works ^3,5,7–12^ have consistently reported substantial intra- and inter-individual variability of TBS-induced neuromodulation ^3^. The variability of TBS outcomes as well as their reproducibility have been mainly assessed with MEPs. MEP amplitude is a compound measure that provides general quantification of both cortical and spinal excitability ^13,14^. TBS-induced changes of MEPs reflect modulation of excitability across the corticospinal tract. However, cortical effects of TBS might not be effectively captured by MEPs, and their validity for drawing conclusions about cortical mechanisms underlying TBS effects is limited.

Resting-state EEG (rsEEG) provides an easy-to-implement noninvasive way to assess spontaneous cortical brain activity both at baseline, and in response to TBS. Multiple studies ^15–20^ have reported associations between MEP amplitudes and the power, as well as phase, of preceding cortical oscillations recorded using EEG. Corticospinal excitability has been frequently linked to EEG activity in alpha and beta frequency bands ^21^. However, studies exploring rsEEG changes following TBS of M1 are scarce and have demonstrated inconsistent results. A single study ^22^ showed that iTBS increases EEG power across a wide range of frequencies (4-90 Hz). In contrast, various studies have reported that cTBS increased EEG power in the theta band (4–7.5 Hz) ^23^, decreased it in delta (2–4 Hz) ^24^ and beta 2 (20–39.5 Hz) ^23^ bands, or produced no change to spectral power ^25^. More importantly, to date only a single iTBS study ^22^ included repeated sessions to assess within subject consistency of the effects. As such, the reliability of TBS-induced modulation of EEG spectral power has not been fully understood.

To address this knowledge gap, using a sham-controlled test-retest design we assessed the reliability of EEG spectral power modulation induced by iTBS and cTBS in 24 healthy participants (aged 18 to 49 years and not taking any psychoactive medications). We increased robustness of our study by including both hypothesis-driven ROI-based and data-driven whole-brain analysis approaches. The ROI-based analysis was focused on EEG signals recorded near the stimulated site within M1, in alpha and beta bands, replicating the methodology already used in earlier studies ^23–25^, and the homologous region in the right hemisphere. Using a ROI over M1 ensures that the results reflect modulation of motor-related brain activity. We hypothesized that TBS would induce modulation of cortical oscillations primarily within the alpha and beta bands in the M1 ROI, but that these effects would not be reliable across visits. The whole-brain exploratory analysis evaluated EEG power across a wide range of frequencies from delta to gamma and in electrodes covering the entire scalp. In addition, we explored both group-based and individual TBS-effects and relationships between TBS-induced modulation of EEG spectral power and corticospinal excitability.

## Results

### TBS-induced modulation of rsEEG power in the left hemisphere

The first analysis was performed by comparing EEG values obtained immediately after TBS (T0), at 15 min (T15) and 25 min (T25) post-TBS time points with the pre-TBS baseline. Hereinafter we will focus on the results of the analysis of relative power in the region of interest in the left frontocentral area (ROI Left). The results of linear mixed models (LMMs) of relative power revealed no significant main effects of the factor Time in both the initial (V1) and retest (V2) visits (all F-values < 2.40, all p-values > 0.05 (corr), partial eta < 0.15, Supplementary Table S1). Figure 1a depicts group-averaged power spectrum (1-50 Hz) obtained in the ROI Left in baseline (pre-TBS) before administering cTBS, iTBS and sham TBS. Individual subject responses to TBS revealed high variability across subjects in both alpha (8 – 12 Hz) and beta 1 (13 – 21 Hz) frequency bands (Figure 1b). Similar variability of relative power was observed from the ROI Right as well as for absolute power in both ROIs (see Supplementary material). This analysis does not support that cTBS, iTBS or sham TBS significantly modulated rsEEG power locally in alpha or beta bands.

**Figure 1.**
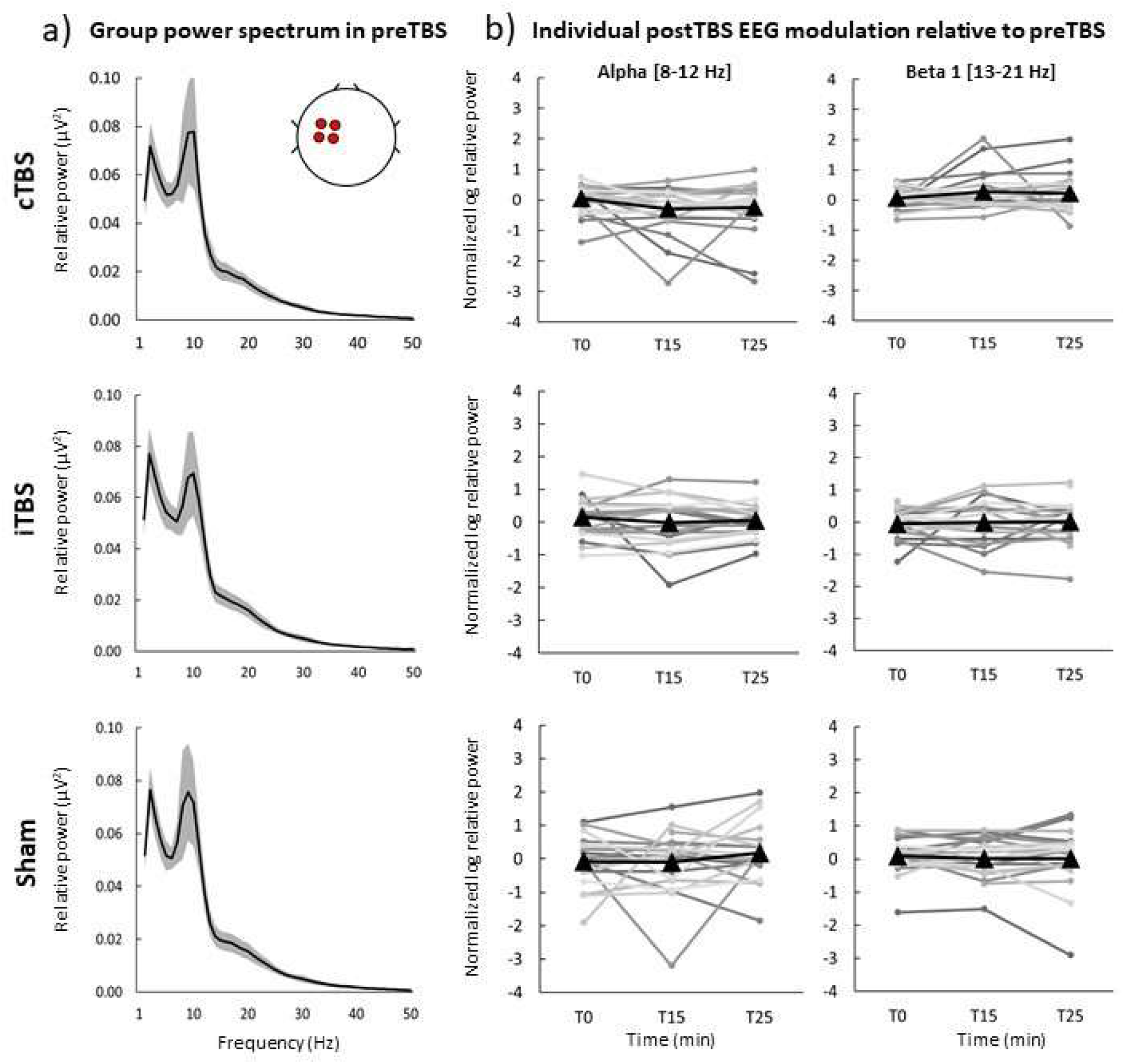
Neuromodulatory effects of cTBS, iTBS and sham TBS on relative power in the initial visit (V1) in ROI Left. a) Group power spectrum (averages across all participants) at 1-50 Hz frequency band for each stimulation protocol in pre-TBS, black line – group mean, grey shaded area – 95% confidence intervals, b) log-transformed ratios post-TBS/pre-TBS in alpha and beta 1 bands for each protocol and post-TBS time point. Black triangles – mean group values for T0, T15 and T25 post-TBS time points, grey round markers – corresponding individual subject values.

### Whole-brain analysis of TBS neuromodulatory effects

To assess neuromodulatory effects of TBS across all electrodes and frequency bands we performed cluster-based permutation analysis of EEG signals in cTBS, iTBS and Sham. We found a significant cluster at T15 in V2 when using the cTBS protocol in the beta and gamma bands (19-43 Hz). This cluster spanned across a broad region of electrodes covering frontal, central, parietal, and occipital areas in both hemispheres (t_max_=924,41.48, p=0.02 (corrected), Figure 2). However, these differences were not observed in V1. There were no other clusters which survived cluster correction. The results of cluster-based permutation analysis of absolute power are presented in the Supplementary material; there were no clusters that survived correction.

**Figure 2.**
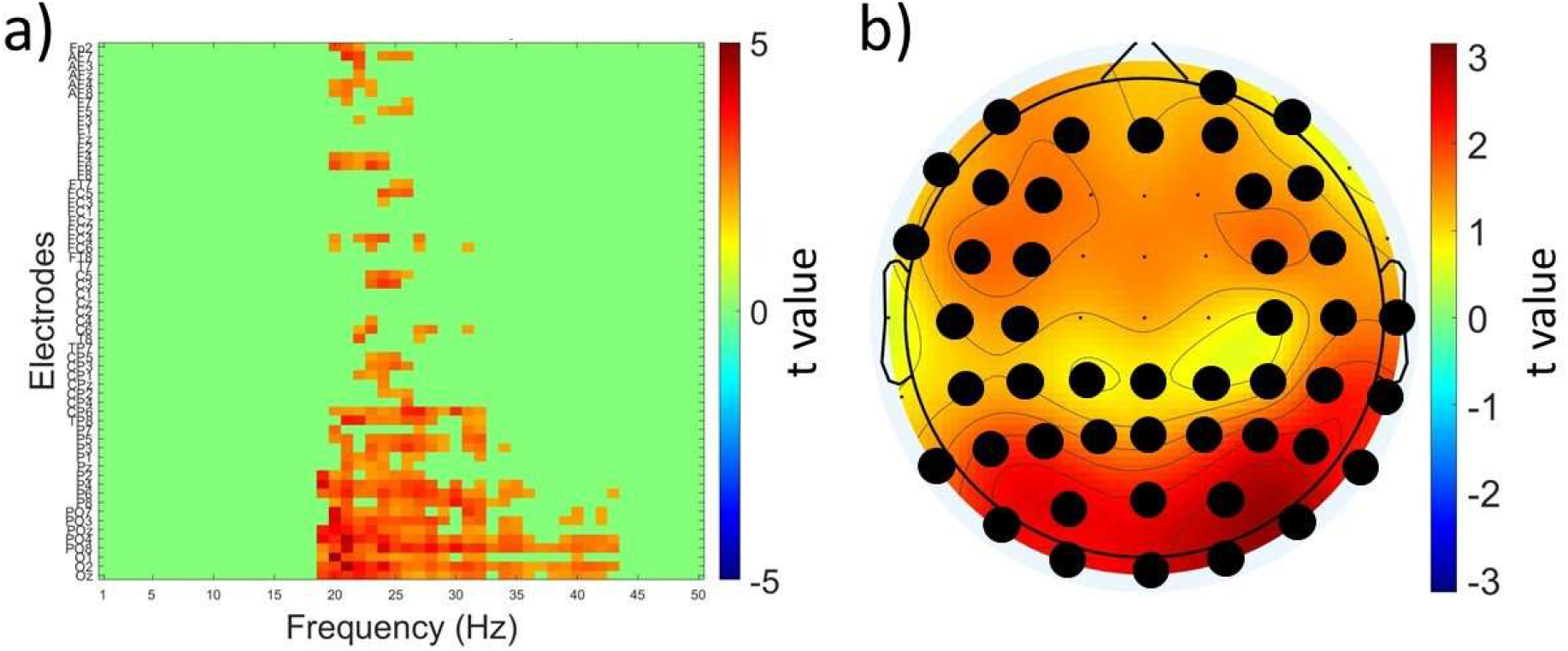
Neuromodulatory effects of cTBS across all electrodes and frequency bands. a) t values associated with whole-brain analysis of relative power using paired t tests comparing T15 and pre-TBS in the retest V2 cTBS visit across 1-50 Hz frequency band (x-axis) and electrodes (y-axis). The color scale is applied only to t-values surviving cluster-based multiple comparisons correction (shown in shades of red), all non-significant values are masked in green; b) topographic representation of the t values averaged across the significant frequencies (19-43 Hz) and electrodes (black dots).

### Reliability of baseline rsEEG power

To address reliability of EEG power across repeated sessions of TBS, we performed a series of reliability analyses. In the first reliability analysis, we compared pre-TBS EEG power in ROI Left across all TBS sessions (Session 1-Ssession 6). The LMMs for relative power yielded no significant main effects of factor Session (all F-values < 1.50, all p-values > 0.05 (corr), partial eta < 0.10, Supplementary Table S3). We found high internal consistency across the six sessions in alpha and beta 1 bands (Cronbach’s α > 0.90, Table S3). This analysis supports that power at baseline did not differ across all stimulation sessions, and the reliability of relative alpha and beta1 power was high. We also found high internal reliability in ROI Right and using absolute power (Table S3).

### Reliability of TBS-induced rsEEG power modulation

We next directly compared TBS-induced power modulation between V1 and V2. To capture change pre to post-TBS, we calculated ratios (post-TBS/pre-TBS) and used them in the analysis input. The results of LMMs of relative power yielded no significant main effect of factor Visit (all F-values < 1.50, all p-values > 0.05 (corr.), partial eta < 0.15, Table S4) with a single exception which did not survive correction for multiple comparisons (Between-visit contrast: iTBS at T0 in alpha band, F=5.25, p=0.03 (uncorrected.), partial eta = 0.20, Table S4). The results of LMMs of relative power in the ROI Right and absolute power in both ROIs are given in the Supplementary material. The results of the test-retest reliability analysis of TBS-induced effects are presented in Figure 3. When using Cronbach’s alpha, we found that reliability of modulation of relative power in alpha and beta 1 bands was mostly low-to-moderate (Cronbach’s α < 0.75). High between-visit reliability was only observed with cTBS at T15 in the alpha band (Cronbach’s α=0.77) and with sham TBS at T25 in the beta 1 band (Cronbach’s α=0.86). The results of LMMs of relative power from the ROI Right and absolute power in both ROIs showed no significant main effect of the factor Visit and low-to-moderate reliability (Supplementary material).

**Figure 3.**
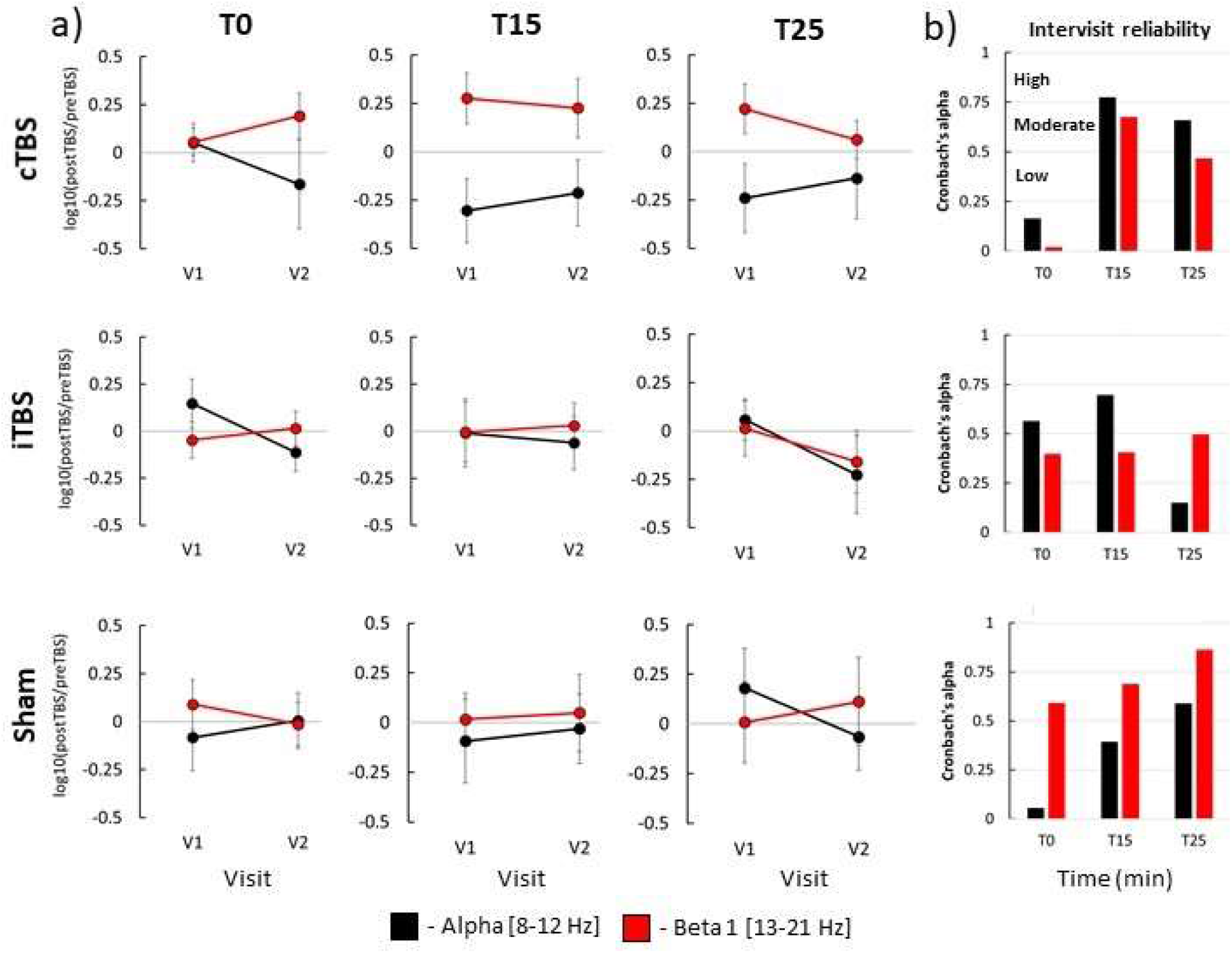
Test-retest reliability of TBS-induced modulation of rsEEG relative power in ROI Left; a) mean post-TBS/pre-TBS ratios at T0, T15 and T25 in V1 and V2 visits in alpha and beta 1 frequency bands. b) Cronbach’s alpha coefficients of consistency between V1 and V2. Spectral power in different frequency bands is shown in different colors: alpha – black, beta 1 – red; error bars - standard error of the mean.

### Contingency analysis of individual TBS-induced changes

As a follow-up analysis to explore characteristics that can be used to classify individual subjects as responders or non-responders to TBS, we analyzed distributions of types of individual TBS-induced rsEEG power changes (increase, decrease, mixed change (increase and decrease at different post-TBS time points), no change) in V1 and V2 in ROI Left in alpha and beta 1 bands. The distributions are presented in Figure 4 a. The proportion of participants who exhibited substantial post-TBS change varied between protocols, visits, and frequency bands but on average was 57±9% (range 45-77%). However, we didn’t observe any clear tendency towards increase or decrease of power attributable to TBS protocol or frequency band. The contingency analysis revealed that distributions of change types were not statistically different between the visits (χ^2^(3, N=43-45)<4.00, p>0.05 (corr.), see Supplementary table S7). We also analyzed the conversion of change types by connecting rsEEG power changes in each participant between the visits. We found that on average 67% of participants exhibited intra-subject variability, with different change types in V1 and V2 (Figure 4, b). Further, both active and sham TBS protocols led to large inter-subject variability of types of EEG changes across the visits (Table S7). Like the results of the analysis of relative power, both active and sham TBS protocols led to large inter-subject variability of types of absolute power changes across the visits (see Supplementary material). In summary, we found no evidence for consistency of TBS response types across subjects and found large intra- and inter-subject variability in response types.

**Figure 4.**
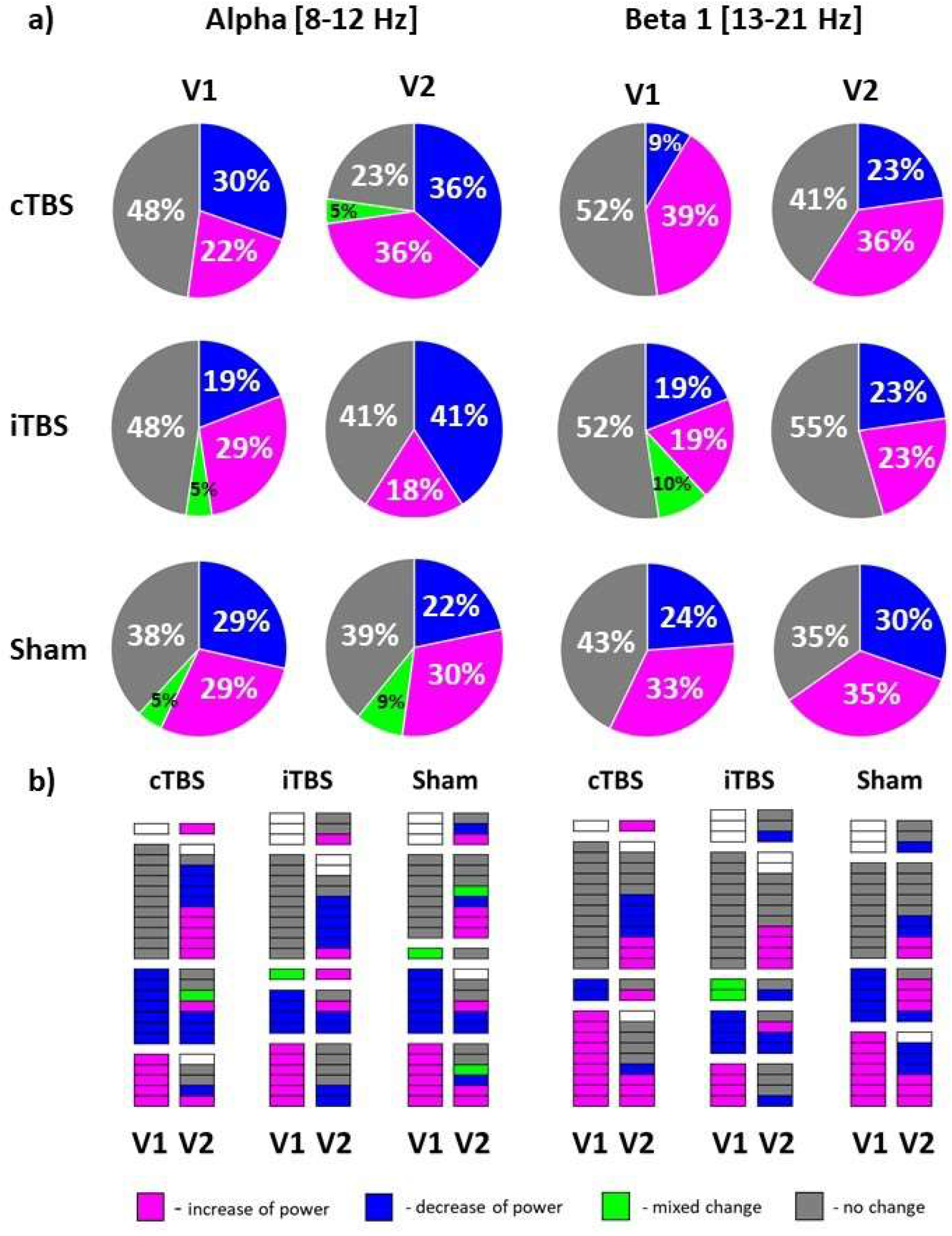
Types of TBS-induced changes of relative EEG power in ROI Left; a) numerical proportions of types in the initial (V1) and retest (V2) visits in alpha and beta 1 bands expressed as percentage; b) conversion plots show types of change in each participant in V1 and V2. Different types of change are given in different colors: magenta - increase of relative power in post-TBS in comparison to pre-TBS, blue – decrease of power, green - mixed change including increase and decrease at different post-TBS time points, grey - no change, white – missing data.

### Comparison of rsEEG power modulation by active and sham TBS protocols

We contrasted neuromodulatory effects between TBS protocols. For this purpose, we conducted separate analyses for each visit and frequency band using power ratios (post-TBS/pre-TBS). The results of LMMs of relative power in ROI Left yielded no significant main effect of the factor Protocol or the Protocol x Time interaction (all F-values < 2.50, all p-values > 0.05 (corr.), partial eta < 0.05, Supplementary table S9) in alpha and beta 1 frequency bands in V1 and V2. These results demonstrate that there were no significant differences between neuromodulatory effects on relative power produced by active and sham TBS protocols in the targeted hemisphere. The results of LMMs of relative power in ROI Right and absolute power yielded a significant main effect of factor Protocol which did not survive correction for multiple comparisons (see Supplementary material).

### Relationship between EEG relative power and corticospinal excitability

Finally, to explore the link between cortical and corticospinal changes following TBS, we analyzed the relationship between modulation of relative rsEEG power in the ROI Left and corticospinal excitability as measured by MEPs. We correlated rsEEG ratios obtained at each post-TBS time point with the MEP data using similar ratios (post-TBS/pre-TBS) collected in the subsequent blocks: rsEEG at T0 with MEPs at T5, rsEEG at T15 with MEPs at T20 and rsEEG and changes of MEPs at T25 with MEPs at T30. We found a significant negative correlation between modulation of relative power in the beta 1 band at T15 and modulation of MEP amplitude at T20 when using iTBS in V1 (Pearson’s r = -0.60, p = 0.01, Figure 5) but this relationship was not replicated in V2 (Pearson’s r = -0.12, p = 0.61). Modulation of relative power in the beta 1 band at T0 positively correlated with MEP changes at T5 when using sham TBS in V1 (Pearson’s r = 0.70, p < 0.01, Figure 5) and in sham V2 (Pearson’s r = 0.54, p = 0.01, Figure 5). There were no other significant correlations between modulation of rsEEG power and MEP amplitudes in the ROI Left. However, modulation of relative power in ROI Right correlated with changes in MEPs in V1 when using iTBS and in both visits when using sham TBS.(see Supplementary material).

**Figure 5.**
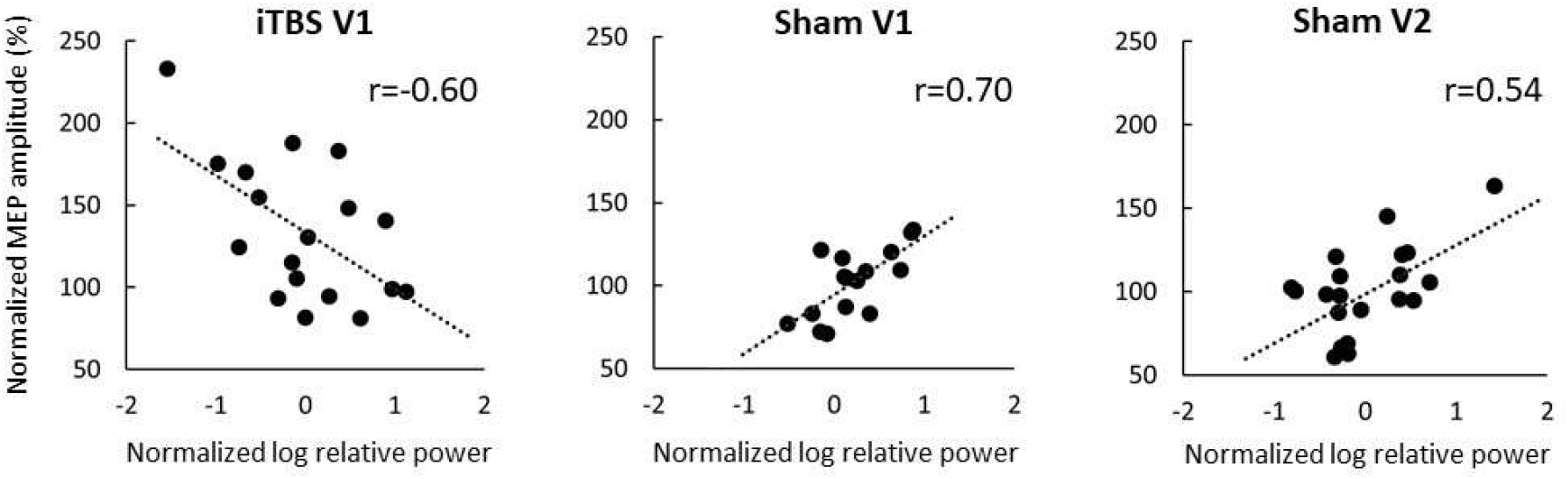
Relationships between modulation of EEG spectral power in beta 1 band and subsequent changes of corticospinal excitability. Scatter plots depicting relationship between normalized log-transformed relative power (post-TBS/pre-TBS ratios) and normalized MEP amplitudes (post-TBS/pre-TBS ratios) in iTBS V1 (EEG at T15 post-TBS and MEP at T20 post-TBS), sham TBS V1 (EEG at T0 post-TBS and MEP at T5 post-TBS), sham TBS V2 (same as in sham V1), dots – individual data points, lines denote linear fit, r - Pearson’s r coefficients.

## Discussion

In this study we assessed the reliability of TBS-induced neuromodulation of resting-state EEG power using a test-retest design with six stimulation sessions (3 protocols with 2 visits for each protocol) for every subject. We explored the effects of cTBS, iTBS, and sham TBS protocols on modulation of spectral EEG power employing a ROI-specific hypothesis-driven analysis in alpha and beta bands, as well as whole-brain data-driven exploratory analyses across a wide range of frequencies (1-50 Hz) and all cap electrodes. We explored the reliability of TBS-induced spectral power modulation at the group level and individual subject level. We also explored whether rsEEG power and MEP amplitudes are co-modulated by TBS.

We found that none of the TBS protocols produced significant group-level effects on ROI-derived rsEEG power. Exploratory whole-brain analysis revealed that cTBS in V2 significantly enhanced relative power in the beta and gamma bands across a broad cortical area; however, these effects were not present in V1. There were no other significant differences between EEG modulation by active and sham TBS protocols. While the reliability of baseline rsEEG power was high across 6 stimulation sessions within individuals, the reliability of TBS-induced neuromodulatory effects was generally low-to-moderate, with substantial inter- and intra-individual variability.

Our group has been systematically investigating the reliability of TBS-induced neuromodulation. This work is a continuation of two previous studies ^12,26^ reporting changes of MEPs and TEPs collected in two consecutive visits in the same subjects as rsEEG analyzed in this report. The results described in the present manuscript using rsEEG are in line with the findings of the earlier studies that TBS effects are highly variable both within and across subjects. Specifically, in our previous works both cTBS and iTBS did not demonstrate consistent modulation of corticospinal excitability, as reflected in MEPs ^12^. Moreover, modulation of early and late components of TEPs were also not reproducible ^26^. Importantly, cTBS and iTBS-induced changes of global and local TMS-evoked cortical responses were not significantly different from sham TBS. In the current study, we also did not find statistically significant differences in modulation of ROI-derived EEG power by active and sham TBS.

The results of the present study support conclusions made in our previous works ^12,26^ that the current knowledge of the mechanisms of TBS-induced neuromodulation, including widely studied “excitatory” or “inhibitory” effects of the iTBS and cTBS protocols, may need to be re-evaluated. For instance, we obtained positive rsEEG power-MEP correlation in beta 1 after sham TBS. A correlation was observed in both visits. Pre-stimulus EEG power in beta band positively correlates with MEP amplitude according to the previous reports ^21^. However, we obtained a negative rsEEG-MEP correlation after active iTBS which was not replicated. Leordori et al., 2021 ^22^ showed that active but not sham iTBS increases EEG power across a wide range (4-90 Hz) of frequencies; however, this effect did not differ between participants with and without MEP facilitation post-TBS. Our data also do not support a linear relationship between TBS modulation of rsEEG power and MEP size. Relationships between TBS modulation of corticospinal excitability and oscillatory brain activity may have a complex nonlinear nature. Moreover, modulation of MEPs and rsEEG power reflect distinct neural mechanisms, such as corticospinal and cortical modulation, that may be contributing to the lack of correlation between them. Further, numerous sources of variability of both rsEEG and MEPs could mask their modulations and relation between them.

Previous studies evaluating TBS modulation of corticospinal ^3,9–14^ and cortical ^26^evoked responses have demonstrated large intra and interindividual variability. Our findings contribute to and expand upon this existing literature by reporting substantial variability of TBS-induced neuromodulatory effects on rsEEG spectral power. Some possible sources of variability across the prior work may be methodological inconsistency and lack of sham control, hindering direct comparison of the results across different studies and assessment of the reproducibility and robustness of observed TBS effects. For example, in a few prior studies, cTBS increased rsEEG power in theta band (4–7.5 Hz) ^23^, decreased it in delta (2–4 Hz) ^24^ and beta 2 (20–39.5 Hz) ^23^ bands, but also produced no change in rsEEG power ^25^. The discrepancy between these results may be explained by the times used between TBS and assessment. The study ^25^ reporting no post-cTBS changes assessed rsEEG modulation only at 5 and 10 minutes, whereas the other studies found significant changes at 20 and 30 minutes post-TBS. Another explanation could be that the studies used data collected during different brain states; one study ^23^reported rsEEG modulation when the subject’s eyes were closed, while the others used data recorded while the subject’s eyes were open. Additionally, with lack of appropriate sham control, it remains unclear how the level of arousal may have influenced the outcomes. There were other methodological differences between the previous studies, including the precise range of the analyzed frequency bands, and location of the electrodes selected for analysis. The previous studies used a small number of selected electrodes for the analysis and the changes of spectral power in these studies were detected close to the stimulation site in the left hemisphere. In contrast, the data-driven approach used in this study may have some methodological advantages over ROI-based approaches in detection of EEG modulation. To our knowledge, there also are no prior studies reporting the reliability of rsEEG modulation within subjects across visits by several TBS protocols. Standardization of experimental procedures and inclusion of appropriate sham condition might improve future research and enable more robust comparisons.

The exact mechanisms underlying TBS effects remain poorly understood. There is evidence suggesting cortical origin of the TBS-induced plasticity and engagement of different physiological mechanisms depending on the type of TBS protocol ^27^. Putative mechanisms of TBS action might include differential modulation of early and late I-waves ^28^, intracortical inhibition ^1,29^, and mediation of neural excitability via N-methyl-D-aspartate receptors involved in LTP/LTD processes ^30^. Another potential mechanism is modulation of rsEEG power through interactions of TBS trains and momentary brain states reflected in ongoing brain oscillations. Understanding of these interactions might be refined by analysis of TBS-induced changes in different frequency bands. This kind of analysis could help to explain the lack of TBS effects on MEPs observed in previous studies and lead to improvements in TBS methodology. In rodent models, it has been shown that during theta oscillations, synapses are in a state of increased plasticity and that synchronizing electrical stimulation to the phase of theta waves enhances LTP induction ^31,32^. However, modulation of rsEEG power in theta band by noninvasive TBS was found in only one out of four previous studies and not in the present work with the whole-brain cluster permutation analysis. Another potential mechanism involving rsEEG power is induction of transient periods of EEG synchronization in the beta band, which is a region-specific intrinsic oscillation in M1 that can be influenced by TMS ^33^. These changes in the beta band were reported in the earlier studies ^22,23^. We also found power modulation in the beta band after cTBS, but it was not reproducible across the visits. Synchronization of TBS bursts with EEG oscillations of various frequencies e.g., individual peak frequency of the sensorimotor mu rhythm may potentially contribute to the stimulation outcomes. In this study we observed changes of absolute rsEEG power in the alpha band in some individuals, but they were not statistically significant at the group level. Importantly, in all studies examining TBS effects on rsEEG, TBS was delivered without synchronization between TMS pulses and endogenous EEG phase and methodological advances in this area might substantially decrease variability of TBS effects on rsEEG.

We note several limitations of the present study. We delivered TMS to an M1 target and therefore, our results cannot be extended to non-motor regions. The lack of reliable TBS effects on conventional outcome metrics of corticospinal excitability (MEPs) limits our interpretation of the rsEEG neuromodulatory effects. Another methodological limitation of this study is that we applied TBS as a percentage of each subject’s active motor threshold (AMT). This is the conventional approach, which enables direct comparison of our results with most of the previous studies. However, the individual stimulation dose was estimated for the excitability level corresponding to voluntary muscle contraction and differences in the relationship of AMT to RMT across subjects were not considered ^34^. Suboptimal stimulation intensity interacting with oscillatory brain activity could potentially invert the sign of TBS-induced changes of spectral power or cancel them totally. Advances in threshold estimation may, therefore, increase the effectiveness of TBS protocols. Our report is limited to the effects of short TBS sessions, whereas longer applications including multiple stimulation sessions might be needed to induce detectable changes of EEG spectral power.

Current experimental evidence suggests that improvement of our understanding of putative neurophysiological mechanisms underlying TBS neuromodulation is needed to achieve consistent TBS effects, which warrants future systematic and rigorous research. We suggest several potential areas for future studies. 1) Standardization of experimental design and TBS protocols is required to draw conclusions about group-level TBS-induced modulations, 2) EEG recordings may be useful for monitoring and adjustment of stimulation parameters to drive cortical TBS effects in the desired direction. 3) Personalizing TBS protocols by adjusting stimulation frequency to the individual peak frequency or region-specific intrinsic oscillations might enable engagement of the corresponding neural mechanisms. 4) Delivering TBS bursts during particular brain states with closed-loop stimulation ^35,36^ appears to be a promising approach to improve TBS reliability. Feasibility of this experimental design has already been demonstrated for MEP outcomes. Targeting with rTMS high-excitability states corresponding to positive peaks of the endogenous sensorimotor µ-rhythm in humans produced LTP-like effect reflected in elevated MEPs ^37^. 5) Accounting for global regulatory mechanisms of plasticity thresholds such as homeostatic metaplasticity could also influence reproducibility of TBS-induced cortical effects. Various priming stimulation protocols, including a priming session followed by a stimulation session have been reported as an effective tool for harnessing metaplastic mechanisms in healthy adults and consequently potentiating the stimulation effects ^38^.

In summary, rsEEG is a noninvasive method to assess modulation of brain activity in response to TBS. This study for the first time systematically investigated effects of active iTBS and cTBS protocols on resting state EEG in the same group of participants and reports no reliable modulation of EEG spectral power using these protocols. We found no significant differences between active and sham TBS. ROI-based and whole-brain analyses did not reveal reproducible rsEEG power changes. Intra- and inter-subject reliability was low-to-moderate. To improve consistency of TBS effects, we suggest that future research should deepen understanding of neural mechanisms underlying TBS effects in different frequency bands and should include standardized methodological practices including appropriate sham control.

## Methods

### Participants

Twenty four healthy volunteers participated in the study (16 males and 8 females, mean age ± SD 30 ± 11 years, range 18 – 49 years). All participants were right-handed (modified Edinburgh handedness inventory ^39^.) None of the participants had contraindications to TMS or magnetic resonance imaging (MRI) ^40,41^, or any history of significant neurological or psychiatric illnesses. Of note, the participants were also not taking any psychoactive medication at the time of measurements. All participants gave their written informed consent to the study. The procedures were approved by the Institutional Review Board of the Beth Israel Deaconess Medical Center, Boston, MA and were in accordance with the Declaration of Helsinki. All procedures were tolerated well by the participants. No unexpected or significant side effects were reported by participants or noticed by experimenters.

### Experimental design

Each volunteer completed two identical sessions of iTBS, cTBS and sham TBS, with sham cTBS and sham iTBS counterbalanced across participants. Each protocol was applied twice, for six stimulation sessions/visits in total. Three initial TBS sessions were performed on separate days (one stimulation session per visit) with at least 2 days in between to eliminate possible carry-over effects. The order of TBS sessions was randomized between subjects. At least 1 month after the initial 3 visits, each participant completed 3 retest visits receiving the same stimulations in the same order. Visits for a given participant were scheduled approximately at the same time of the day to control for possible circadian influences on the neuromodulatory effects of TMS ^42^. During each visit rsEEG data, MEPs, and TMS-evoked EEG potentials (TEPs) were collected. MEP and TEP results were published previously ^12,26^. Blocks of EEG data in the eyes-open resting state, 2.5-3 minutes in duration, were collected before the administration of one of the TBS protocols (pre-TBS), and then again immediately after TBS, and at 15 and 25 minutes (T0, T15 and T25). EEG data of 3 participants in cTBS condition (1 in V1, 2 in V2), 5 participants in iTBS condition (3 in V1, 2 in V2) and 4 in sham TBS condition (3 in V1, 1 in V2) were not included in the study due to missing or corrupted EEG files, or excessive artifacts. The MEP data that were used in the correlation analysis were collected immediately after each rsEEG block, at 5, 20 and 30 min after TBS (T5, T20 and T30 correspondingly). The experimental design (Fig. S1), details of MRI acquisition, determination of motor hotspot, resting and active motor threshold, and methods used for MEP recordings and analyses are presented in the Supplementary material.

### Real and sham TBS

All TBS protocols were applied to the individual participant’s motor hotspot at 80% of AMT in each participant. The iTBS and cTBS protocols consisted of 600 pulses in total given over 200 s and 40 s time periods correspondingly. The iTBS protocol included trains of 3-pulse bursts (duration of trains 2 s, burst frequency 50 Hz, inter-burst interval 200 ms) spaced by 8 s ITIs. The cTBS protocol used 3 pulse bursts (frequency 50 Hz, inter-burst interval 200 ms) that were delivered continuously. In addition to iTBS and cTBS, each volunteer was randomly assigned to receive either sham iTBS or sham cTBS during the initial visit series, and the same sham stimulation during retest visit series. For administration of the sham TBS protocols, the placebo side of the Cool-B65 A/P coil with a 3D printed 3.3 cm spacer (MagVenture A/S, Farum, Denmark) attached to the placebo side was used. Single positive triangle electrical pulses (pulse width 2.2 ms, intensity 2-3 mA) were given synchronously with TMS and sham TMS for additional blinding of the participants by an attempt to match the sensory experience of the active and sham protocols. For this purpose, a pair of self-adhesive surface electrodes (Ambu Neuroline 715, Ambu A/S Baltorpbakken 13, DK-2750 Ballerup) were placed approximately 1 cm below the inion. The electrical stimulation was applied synchronously with all TBS protocols.

### EEG recording and pre-processing

RsEEG data were collected with a 64-channel TMS-compatible system (actiCHamp, Brain Products GmbH, Munich, Germany) at a sampling rate of 1000 Hz and labeled according to the extended 10–20 international system. The Fp1 electrode was used as an online reference and Fpz was used as a ground. The electrode impedances were monitored throughout the sessions and kept below 5 kΩ during each session. EEG signal digitization and online monitoring were performed with a BrainCHamp DC amplifier and BrainVision Recorder software (version 1.21). EEG recordings were pre-processed offline using EEGLAB toolbox v2021.0 ^43^ and a custom script written in Matlab 2020b (MathWorks, Inc., Natick, MA, United States). High-pass (1 Hz) and low-pass (50 Hz) fourth-order Butterworth filters were applied. Bad EEG channels and noise or artifact bursts were automatically removed using the clean_rawdata EEGLAB plugin v2.3. This procedure resulted in removal of less than 1% of the data. EEG data were re-referenced to the average across channels and divided into 6-second epochs. Two individual datasets (one in cTBS V1 and one in sham TBS V1 condition) were excluded due to the short length of EEG recording resulting in a small (<5) number of epochs. All data across channels were concatenated into a single file for each participant separately to use in the next step of the pre-processing pipeline. EEG data were visually inspected and remaining bad channels and bad epochs were manually removed (total average ±SD automatically and manually rejected channels = 1.6 ±1.3, average ±SD rejected epochs = 2.0 ±4.7, average ±SD remaining epochs = 24.1 ±3.2). Rejected channels were interpolated using spherical interpolation. Dimensionality of EEG data was reduced with principal component analysis (PCA) resulting in 30 dimensions (with exception of 35 dimensions in one dataset and 40 dimensions in 11 datasets with lower signal quality) prior to independent component analysis. This procedure can improve signal to noise ratio and decomposition of large sources ^44^. Fast ICA (fICA v2.5) ^45^ and Multiple Artifact Rejection Algorithm (MARA v1.2) ^46^ EEGLAB plugins were used to compute, classify, and automatically remove independent components (ICs) corresponding to blink/eye movement, electromyographic activity, single electrode noise, or cardiac beats artifacts. Remaining artifact ICs mainly containing muscle activity were manually removed using the TMS-EEG Signal Analyzer (TESA v1.1.1) EEGLAB toolbox ^47^ (total number of analyzed ICs was 30 for most subjects (35 - 40 ICs in 12 datasets with lower signal quality), total average ±SD rejected ICs was 18.3 ±5.3; total average ±SD remaining ICs was 12.6 ±3.8). Cleaned EEG data were unmerged into pre-TBS, T0, T15 and T25 files.

### EEG analyses

#### ROI analysis

Absolute EEG power was computed for the 1 - 50 Hz frequency band for each electrode using the spectopo EEGLAB function (window-size - 1000 samples, window-overlap - 500 samples). Absolute and relative EEG power were calculated for two regions of interest (ROI Left and ROI Right) and two frequency bands: alpha (8-12 Hz) and beta1 (13-21 Hz). Absolute power was computed by summing ROI-averaged power across all frequency bins in each band. Relative EEG power was obtained by dividing the absolute power values in each frequency bin by the sum of absolute power values across the 1 - 50 Hz frequency band. ROI Left included four electrodes (FC3 FC1 C3 C1) surrounding the average stimulation site in the left M1. ROI Right included four homologous electrodes (FC2 FC4 C2 C4) located over the contralateral area in the right hemisphere. Statistical analyses were performed using the JMP Pro version 16.1.0 (SAS Institute Inc., Cary, NC, USA) and Jamovi version 2.3.2.0. (The jamovi project (2021), https://www.jamovi.org, Sydney, Australia). All data were checked for normality using the Shapiro-Wilk test and log10-transformed. To evaluate neuromodulatory effects of each TBS protocol on rsEEG in alpha and beta1 bands in V1 and V2, ROI-averaged absolute and relative power values for each protocol, visit, frequency band and ROI were entered into separate LMMs with Time (pre-TBS, T0, T15, T25) as the independent variable crossed with a random effect for subjects.

#### Cluster-permutation analysis

In a data-driven analysis, we also assessed TBS-induced EEG modulation across all 59 electrodes and frequencies (1-50 Hz) using a cluster-based permutation analysis. We created a neighborhood matrix to define neighboring electrodes. The spatial neighborhood of each electrode included all electrodes within 4 cm around it (mean 4.2±0.93, range 2-7 neighbors per electrode). The data recorded from the channels FT9, FT10, TP9 and TP10 were contaminated by artifacts in most of the datasets. To ensure a high signal to noise ratio these data were excluded from the analysis. These channels had a single neighboring channel in contrast to other channels and therefore their exclusion did not affect the neighborhood matrix. Pairwise comparisons between pre-TBS and post-TBS time points were calculated using paired-samples t-tests. A nonparametric cluster-based permutation approach was utilized to control for multiple comparisons across electrode-frequency space ^48^. A cluster was defined as at least two neighboring significant (p < 0.05) data points in either frequency or space. After permuting the data 1000 times, clusters with a cluster statistics (sum of t-values included in the cluster) exceeding 97.5% of the respective null distribution cluster statistics were considered significant. The cluster building procedure was performed separately for data points with positive and negative t-values.

#### Baseline rsEEG analysis

To test if baseline measurements differed across the six sessions, pre-TBS absolute and relative power for each frequency band and ROI were entered into separate LMMs with Session (S1, S2, S3, S4, S5, S6) as the independent variable crossed with a random effect for subjects.

*Test-retest (reliability) analysis:* To compare the effects of each TBS protocol between V1 and V2 we used baseline corrected values computed as post-TBS/pre-TBS ratios. Thus, the baseline corrected post-TBS absolute and relative power values for each ROI, frequency band and post-TBS time point were entered into separate LMMs with Visit (V1, V2) as the independent variable crossed with random effect for subjects. For assessment of test–retest reliability of baseline EEG measures and TBS-induced effects, Cronbach’s α coefficients were computed between the visits for absolute and relative power in each ROI and frequency band separately. Individual TBS-induced EEG changes were analyzed by comparison of EEG power values computed in each epoch in pre-TBS and post-TBS using Wilcoxon’s signed rank test. Thus, individual EEG changes in each participant were classified into four categories: increase – post-TBS power values were significantly larger than pre-TBS in at least one post-TBS time point, decrease – post-TBS power values were significantly smaller than pre-TBS in at least one post-TBS time point, mixed change – post-TBS power values were significantly larger and smaller than pre-TBS in different post-TBS time points, no change – absence of significant post-TBS-pre-TBS differences.

#### Sham-controlled analysis

To compare TBS effects between protocols post-TBS/pre-TBS ratios of absolute and relative power values for each ROI, frequency band and time point were entered into separate LMMs with Protocol (cTBS, iTBS, Sham) as the independent variable and an interaction of Protocol x Time crossed with a random effect for subjects.

#### rsEEG-MEP correlations

To assess the relationships between TBS-induced modulation of rsEEG power and corticospinal excitability Pearson’s correlations between rsEEG ratios obtained at each post-TBS time point with the MEP ratios collected in the following blocks were used (EEG at T0 with MEPs at T5, EEG at T15 with MEPs at T20 and EEG at T25 with MEPs at T30). Bonferroni correction for multiple comparisons was applied where appropriate. The data are given as mean ± standard deviation (SD).

## Acknowledgements

We are grateful to all research assistants who helped to run these study visits. We are grateful to the gracious funding from the MIT Harvard Broad institute (6600024e5500000895) directly supporting the study and our line of research on brain plasticity biomarkers. Dr. Santarnecchi is supported by the Defence Advanced Research Projects Agency (DARPA) via HR001117S0030, the NIH (P01 AG031720-06A1, R01 MH117063-01, R01 AG060981-01) and ADDF (ADDF-FTD GA201902e2017902). Dr. Pascual-Leone is supported by grants from the National Institutes of Health (R24AG06142, and P01 AG031720), the National Science Foundation, and the Barcelona Brain Health Initiative funded primarily by La Caixa. Dr. Shafi is supported by the Football Players Health Study (FPHS) at Harvard University, and the NIH (R01MH115949, R01AG060987, P01 AG031720-06A1). Dr. Benwell is supported by the British Academy/Leverhulme Trust and the United Kingdom Department for Business, Energy and Industrial Strategy [SRG19/191169]. During manuscript preparation, Dr. Ross was supported by the Department of Veterans Affairs Office of Academic Affiliations Advanced Fellowship Program in Mental Illness Research and Treatment, the Medical Research Service of the Veterans Affairs Palo Alto Health Care System and the Department of Veterans Affairs Sierra-Pacific Data Science Fellowship. The content of this paper is solely the responsibility of the authors and does not necessarily represent the official views of Harvard University and its affiliated academic health care centers, or the National Institutes of Health.

## Author contributions

E.S., A.P.L., M.S. conceptualized the study. P.F., D.M., R.O. and P.B contributed to specific methodological aspects. P.B., R.O., and D.M. ran the study visits and collected the data. A.R. preprocessed and analyzed the data with contributions from R.O, C.B., P.F. J.R. and M.S. A.R. wrote the original draft of the manuscript with general oversight by M.S. All authors reviewed the manuscript.

