## Supplementary material for "Reliability of resting-state EEG modulation by continuous and intermittent theta burst stimulation of the primary motor cortex: A sham-controlled study"

### Supplementary methods

#### MRI acquisition

A T1-weighted structural MR image was obtained for each subject with a 3T MRI scanner (GE Healthcare, Ltd., United Kingdom) and used for TMS navigation. MRIs were taken using a 3D spoiled gradient echo sequence (166 axial slices, 240-mm isotropic field-of view, voxel size 0.937 × 0.937 × 1 mm, flip angle = 15°, TE/TR ≥ 2.9/6.9 ms, duration ≥ 432 s).

#### Determination of motor hotspot, resting and active motor threshold

Participants were comfortably seated in an armchair and were instructed to keep their eyes open and their muscles relaxed during the experiments. Monophasic single-pulse TMS (spTMS) was delivered to the left primary motor cortex (M1) with a MagPro Cool-B65 A/P figure-of-eight coil (75mm outer wing diameter) and a MagPro X100 stimulator (MagVenture A/S, Farum, Denmark). Brainsight™ TMS Frameless Navigation system (Rogue Research Inc., Montreal, Canada) was used for real time monitoring of accurate and reproducible coil positioning within and across sessions. Motor hotspot, resting and active motor thresholds (RMT and AMT) were found in each participant for the first dorsal interosseous (FDI) muscle. For motor hotspot search the coil was placed over the hand area of M1 and oriented to induce a posterior–anterior current perpendicular to the central sulcus. The hotspot was defined as the location within M1 where TMS induced the largest and consistent MEPs. RMT was defined as the lowest stimulator intensity expressed as percentage of maximum stimulator output (%MSO) to elicit MEPs of at least 50 µV peak-to-peak amplitude in 5 out of 10 consecutive trials in relaxed target muscle. AMT was defined as the lowest %MSO to elicit MEPs of at least 200 µV in 5 out of 10 consecutive trials in the voluntary contracted target muscle. Participants wore earplugs throughout the stimulation sessions for hearing protection.

#### Motor evoked potentials (MEPs)

Motor evoked potentials (MEPs) were recorded from the first dorsal interosseous (FDI) muscle of the right hand. In pre-TBS 120 single-pulse TMS (spTMS) were given and corresponding MEPs were recorded. After that, one of three TBS protocols was applied, followed by blocks of 60 spTMS. For the purpose of this study we selected and analyzed only MEPs recorded within the blocks at 5, 20 and 30 min after the TBS. Mean MEP amplitudes for each post-TBS time-point were expressed as the percentage change from pre-TBS. For details of MEP analysis see the previous study [1].

### Supplementary results

**TBS-induced modulation of EEG relative power in the right hemisphere and absolute power in both hemispheres**

The results of LMMs of relative power on ROI Right revealed no significant main effects of factor Time in V1 and V2 (all F-values < 3.00, all p-values > 0.05 (corr), partial eta < 0.15, Supplementary Table S1). The results of LMMs of absolute power on ROI Left and Right revealed a single main effect of factor Time in the ROI Left in cTBS V2 in beta 1 band which didn't survive correction for multiple comparison (F-value = 2.76, p-values > 0.05 (corr), partial eta < 0.12, Supplementary Table S2).

**Whole-brain analysis of TBS neuromodulatory effects on EEG absolute power**

The analysis of absolute power didn't identify any significant clusters.

**Modulation of corticospinal excitability**

We briefly describe the results of MEP analysis previously published by [1]. MEP amplitudes were increased by iTBS and sham TBS in V1 and V2 (Figure S2). cTBS had facilitatory effect on MEPs in V2. For details of results of MEP analysis see the previous study [1].

**Comparisons of EEG modulation between visits and test-retest reliability of TBS effects**

The results of LMM of relative power in the ROI Right yielded no significant main effect of factor Visit (all F-values < 3.00, all p-values > 0.05 (corr.), partial eta < 0.15, Table S4). The results of LMM of absolute power in the ROI Left and ROI Right yielded no significant main effect of factor Visit (all F-values < 2.00, all p-values > 0.05 (corr.), partial eta < 0.10, Table S5). Overall, reliability of modulation of relative power in ROI Right and absolute power in both ROIs was low-to-moderate (Cronbach's a < 0.75, Table S6). High between-visit reliability of modulation of relative power was obtained in the ROI Right in cTBS at T15 in beta 1 band (Cronbach's a=0.83, Table S6) and in sham TBS also in beta 1 band at T15 (Cronbach's a=0.78, Table S6) and T25 (Cronbach's a=0.98, Table S6). Modulation of absolute EEG power was highly consistent between two visits in the ROI Right after sham TBS in alpha band at T25 (Cronbach's a=0.85, Table S6).

**Contingency analysis of individual TBS-induced changes**

The contingency analysis revealed that distributions of change types were not statistically different between the visits ($\chi^2$(3, N=43-45)<7.00, p>0.05 (corr.), see Supplementary table S7). The amount of participants who exhibited significant post-TBS change varied between protocols, visits and frequency bands being on average 76±13% (range 52 and 95%). There was no any clear tendency towards increase or decrease of power attributable to TBS protocol or frequency band. On average 65% of participants exhibited different change types in V1 and V2 (Figure S3). Like the results of analysis of relative power, both active and sham TBS protocols led to large inter-subject variability of types of EEG changes across the visits (cTBS – 64±10%, iTBS – 79±0%, sham TBS – 53±18%, average across alpha and beta1 bands).

**Comparison of EEG modulation by active and sham TBS protocols**

The results of LMMs of relative power in ROI Right yielded significant main effect of factor Protocol in beta 1 band in V1 which didn't survive correction for multiple comparison (F-value = 4.69, p-values > 0.05 (corr), partial eta = 0.05). The results of LMMs of absolute power in ROI Left and ROI Right also yielded significant main effects of factor Protocol in beta 1 band in V2 which didn't survive correction for multiple comparison (F-value = 4.81, p-values > 0.05 (corr), partial eta = 0.05, and F-value = 3.31, p-values > 0.05 (corr), partial eta = 0.04 correspondingly).

**Relationship between EEG spectral power and corticospinal excitability**

We performed analysis of relationships between modulation of EEG power in alpha and beta 1 bands in the non-targeted right hemisphere (ROI Right) and corticospinal excitability of the contralateral side by correlating EEG changes and MEPs in those protocols and time points where correlations in the targeted left hemisphere (ROI Left) were found. There was a negative correlation between modulation of relative power in the ROI Right in beta 1 band at T15 and MEP modulation in the left hemisphere at T20 in iTBS V1 (Pearson's r = -0.52, p < 0.05). Modulation of relative power in beta 1 band at T0 in the ROI Right positively correlated with MEP changes in the left hemisphere at T5 in sham V1 (Pearson's $r = 0.60$, $p < 0.05$) and in sham V2 (Pearson's $r = 0.53$, $p = 0.01$).We also found positive correlation between modulation of absolute EEG power in the ROI Right in alpha band at T15 and MEP modulation at T20 in iTBS V1 (Pearson's $r = 0.48$, $p < 0.05$).

### Supplementary figures and tables

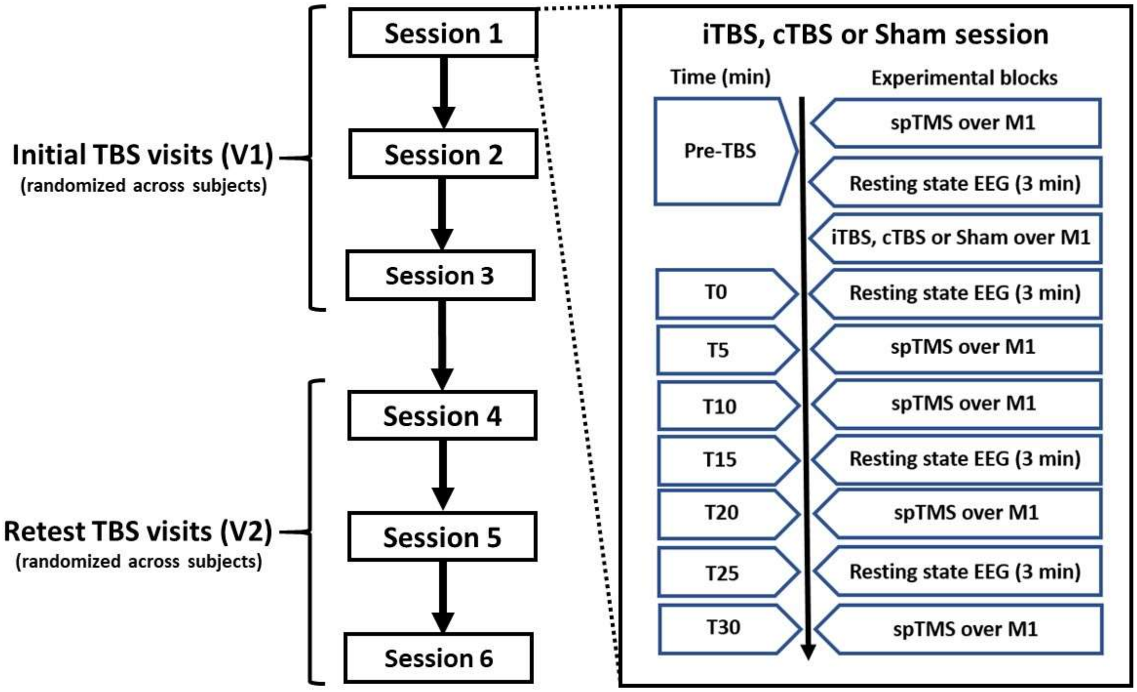


Supplementary Figure S1. Experimental design. spTMS – single pulse TMS, M1 – the primary motor cortex

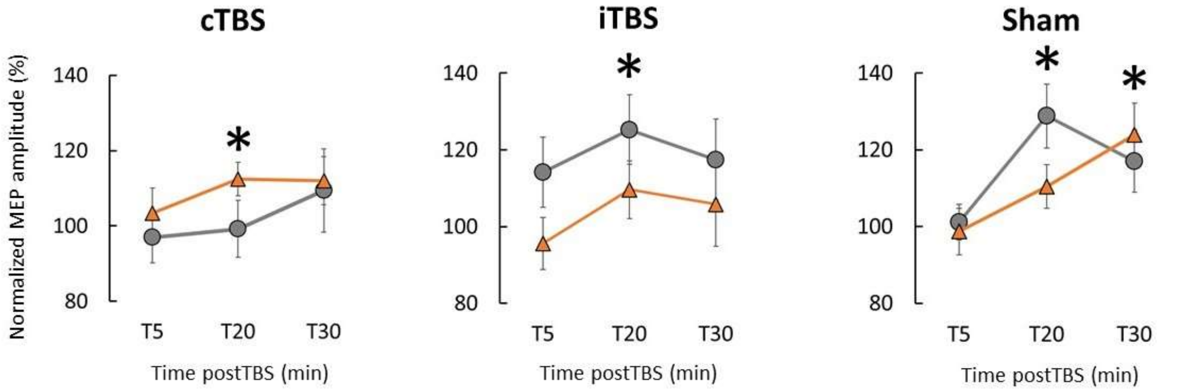


Supplementary Figure S2. Modulation of corticospinal excitability by cTBS, iTBS and sham TBS. Gray round markers - group means of normalized MEP amplitudes (post-TBS/pre-TBS ratios) at 5, 20 and 30 min post-TBS in the initial visit (V1), orange triangles - corresponding values in the retest visit (V2). Error bars – standard error of the mean. Asterisks denote significant MEP changes in comparison to 100% pre-TBS ($p < 0.05$). Significant MEP facilitation by iTBS in V2 occurred at T10 (not shown).

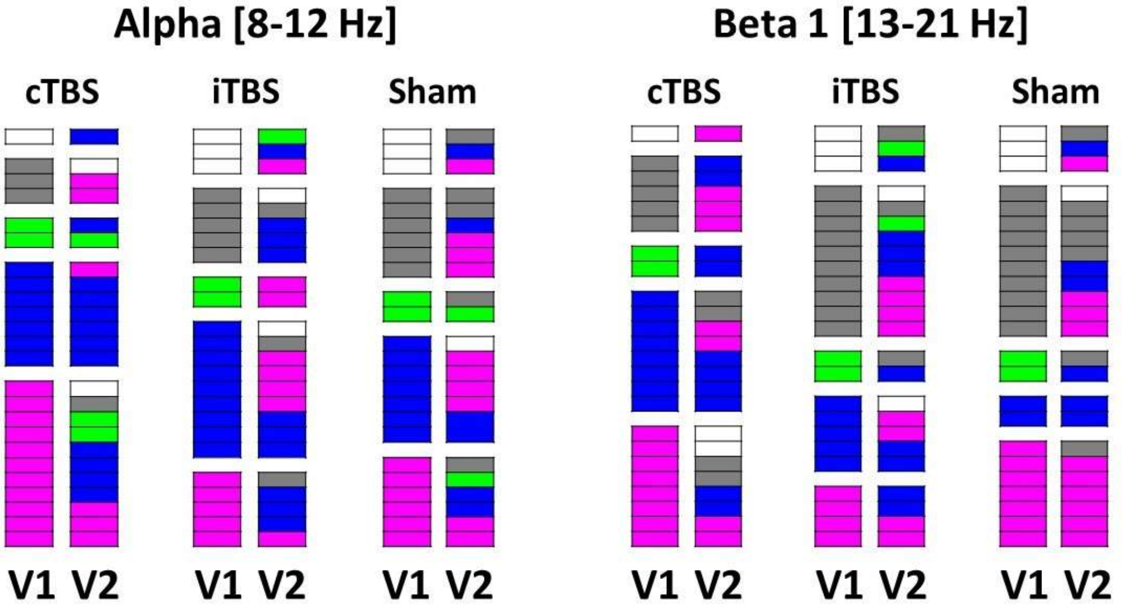


Supplementary figure S3. Conversion of EEG change types in each participant in alpha and beta 1 bands in the initial (V1) and retest (V2) visits. Different types of change are given in different colors: magenta - increase of absolute power in post-TBS in comparison to pre-TBS, blue – decrease of power, green - mixed change including increase and decrease at different post-TBS time points, grey - no change, white – missing data.

Supplementary Table S1. Results of linear mixed model analysis of TBS neuromodulatory effects on relative EEG power in ROI Left and ROI Right in the initial (V1) and retest (V2) visits

| Protocol | Visit | Frequency | ROI Left | | | | | | ROI Right | | | | | |
|---|---|---|---|---|---|---|---|---|---|---|---|---|---|---|
| | | | N parm | DF | DF Den | F-value | p-value (uncorr) | Partial eta | N parm | DF | DF Den | F-value | p-value (uncorr) | Partial eta |
| cTBS | V1 | Alpha | 3 | 3 | 65.02 | 2.322 | 0.083 | 0.10 | 3 | 3 | 65.04 | 1.737 | 0.168 | 0.07 |
| | | Beta 1 | 3 | 3 | 65.02 | 1.953 | 0.130 | 0.08 | 3 | 3 | 65.03 | 2.635 | 0.057 | 0.11 |
| | V2 | Alpha | 3 | 3 | 63.00 | 0.415 | 0.743 | 0.02 | 3 | 3 | 63.00 | 0.327 | 0.806 | 0.02 |
| | | Beta 1 | 3 | 3 | 63.00 | 1.249 | 0.299 | 0.06 | 3 | 3 | 63.00 | 0.817 | 0.489 | 0.04 |
| iTBS | V1 | Alpha | 3 | 3 | 58.04 | 0.497 | 0.686 | 0.03 | 3 | 3 | 58.04 | 0.200 | 0.896 | 0.01 |
| | | Beta 1 | 3 | 3 | 58.05 | 0.102 | 0.959 | 0.01 | 3 | 3 | 58.05 | 0.157 | 0.925 | 0.01 |
| | V2 | Alpha | 3 | 3 | 65.03 | 0.786 | 0.506 | 0.04 | 3 | 3 | 65.03 | 0.425 | 0.736 | 0.02 |
| | | Beta 1 | 3 | 3 | 65.02 | 0.887 | 0.452 | 0.04 | 3 | 3 | 65.03 | 1.022 | 0.389 | 0.05 |
| Sham | V1 | Alpha | 3 | 3 | 57.16 | 0.839 | 0.478 | 0.04 | 3 | 3 | 57.13 | 0.782 | 0.509 | 0.04 |
| | | Beta 1 | 3 | 3 | 57.03 | 0.048 | 0.986 | 0.00 | 3 | 3 | 57.03 | 0.242 | 0.867 | 0.01 |
| | V2 | Alpha | 3 | 3 | 64.00 | 0.092 | 0.964 | 0.00 | 3 | 3 | 64.00 | 0.080 | 0.971 | 0.00 |
| | | Beta 1 | 3 | 3 | 64.10 | 0.259 | 0.854 | 0.01 | 3 | 3 | 64.10 | 0.049 | 0.986 | 0.00 |

DF – degrees of freedom for the numerator, DF Den – degrees of freedom for the denominator, p-value (uncorr) – uncorrected p-value, ROI Left and ROI Right – regions of interest in the left and right hemisphere

Supplementary Table S2. Results of linear mixed model analysis of TBS neuromodulatory effects on absolute EEG power in the initial (V1) and retest (V2) visits

| Protocol | Session | Frequency | ROI Left | | | | | | ROI Right | | | | | |
|---|---|---|---|---|---|---|---|---|---|---|---|---|---|---|
| | | | N parm | DF | DF Den | F-value | p-value (uncorr) | Partial eta | N parm | DF | DF Den | F-value | p-value (uncorr) | Partial eta |
| cTBS | V1 | Alpha | 3 | 3 | 65.00 | 1.561 | 0.207 | 0.07 | 3 | 3 | 65.03 | 1.490 | 0.225 | 0.06 |
| | | Beta 1 | 3 | 3 | 65.00 | 0.692 | 0.560 | 0.03 | 3 | 3 | 65.01 | 0.751 | 0.526 | 0.03 |
| | V2 | Alpha | 3 | 3 | 63.00 | 1.977 | 0.127 | 0.09 | 3 | 3 | 63.00 | 1.400 | 0.251 | 0.06 |
| | | Beta 1 | 3 | 3 | 63.00 | 2.757 | **0.049** | 0.12 | 3 | 3 | 63.00 | 2.244 | 0.092 | 0.10 |
| iTBS | V1 | Alpha | 3 | 3 | 58.02 | 0.056 | 0.983 | 0.00 | 3 | 3 | 58.01 | 0.132 | 0.941 | 0.01 |
| | | Beta 1 | 3 | 3 | 58.00 | 0.520 | 0.670 | 0.03 | 3 | 3 | 58.00 | 0.863 | 0.466 | 0.04 |
| | V2 | Alpha | 3 | 3 | 65.03 | 0.217 | 0.885 | 0.01 | 3 | 3 | 65.02 | 0.123 | 0.946 | 0.01 |
| | | Beta 1 | 3 | 3 | 65.02 | 0.209 | 0.890 | 0.01 | 3 | 3 | 65.01 | 0.127 | 0.944 | 0.01 |
| Sham | V1 | Alpha | 3 | 3 | 57.06 | 1.046 | 0.379 | 0.05 | 3 | 3 | 57.06 | 0.929 | 0.433 | 0.05 |
| | | Beta 1 | 3 | 3 | 57.01 | 1.176 | 0.327 | 0.06 | 3 | 3 | 57.00 | 1.330 | 0.274 | 0.07 |
| | V2 | Alpha | 3 | 3 | 64.08 | 0.087 | 0.967 | 0.00 | 3 | 3 | 64.08 | 0.087 | 0.967 | 0.00 |
| | | Beta 1 | 3 | 3 | 64.04 | 0.923 | 0.435 | 0.04 | 3 | 3 | 64.03 | 0.506 | 0.679 | 0.02 |

DF – degrees of freedom for the numerator, DF Den – degrees of freedom for the denominator, p-value (uncorr) – uncorrected p-value, ROI Left and ROI Right – region of interest in the left and right hemisphere, bold font – significant p-value before correction.

Supplementary Table S3 Results of linear mixed model and reliability analysis of relative and absolute EEG power at the baseline (pre-TBS) in the ROI Left and Right across six stimulation sessions

| Frequency | ROI Left | | | | | | | ROI Right | | | | | | |
|---|---|---|---|---|---|---|---|---|---|---|---|---|---|---|
| | N parm | DF | DF Den | F-value | p-value (uncorr) | Partial eta | Cronbach's alpha | N parm | DF | DF Den | F-value | p-value (uncorr) | Partial eta | Cronbach's alpha |
| | Relative power | | | | | | | Relative power | | | | | | |
| Alpha | 5 | 5 | 104.2 | 1.033 | 0.402 | 0.05 | 0.98 | 5 | 5 | 104.2 | 0.348 | 0.883 | 0.02 | 0.97 |
| Beta 1 | 5 | 5 | 104.1 | 0.292 | 0.917 | 0.01 | 0.99 | 5 | 5 | 104.1 | 0.202 | 0.961 | 0.01 | 0.99 |
| | Absolute power | | | | | | | Absolute power | | | | | | |
| Alpha | 5 | 5 | 104.1 | 1.353 | 0.248 | 0.06 | 0.98 | 5 | 5 | 104.1 | 0.374 | 0.866 | 0.02 | 0.98 |
| Beta 1 | 5 | 5 | 104.1 | 1.454 | 0.211 | 0.07 | 0.98 | 5 | 5 | 104.0 | 0.318 | 0.901 | 0.02 | 0.98 |

DF – degrees of freedom for the numerator, DF Den – degrees of freedom for the denominator, p-value (uncorr) – uncorrected p-value, ROI Left and ROI Right – region of interest in the left and right hemisphere

Supplementary table S4. Results of linear mixed model analysis of TBS-induced modulation of relative EEG power in ROI Left and ROI Right across visits

| Protocol | Frequency | Time postTBS | ROI Left | | | | | | ROI Right | | | | | |
|---|---|---|---|---|---|---|---|---|---|---|---|---|---|---|
| | | | N parm | DF | DF Den | F-value | p-value (uncorr) | Partial eta | N parm | DF | DF Den | F-value | p-value (uncorr) | Partial eta |
| cTBS | Alpha | T0 | 1 | 1 | 23.24 | 0.812 | 0.377 | 0.03 | 1 | 1 | 22.84 | 0.127 | 0.725 | 0.01 |
| | | T15 | 1 | 1 | 21.81 | 0.401 | 0.533 | 0.02 | 1 | 1 | 22.30 | 0.469 | 0.501 | 0.02 |
| | | T25 | 1 | 1 | 22.33 | 0.181 | 0.675 | 0.01 | 1 | 1 | 22.49 | 0.079 | 0.782 | 0.00 |
| | Beta 1 | T0 | 1 | 1 | 22.58 | 1.003 | 0.327 | 0.04 | 1 | 1 | 23.36 | 0.013 | 0.910 | 0.00 |
| | | T15 | 1 | 1 | 22.43 | 0.094 | 0.762 | 0.00 | 1 | 1 | 21.64 | 1.908 | 0.181 | 0.08 |
| | | T25 | 1 | 1 | 22.96 | 1.213 | 0.282 | 0.05 | 1 | 1 | 23.23 | 2.671 | 0.116 | 0.10 |
| iTBS | Alpha | T0 | 1 | 1 | 20.41 | 5.251 | **0.033** | 0.20 | 1 | 1 | 21.87 | 0.314 | 0.581 | 0.01 |
| | | T15 | 1 | 1 | 24.49 | 0.016 | 0.902 | 0.00 | 1 | 1 | 24.35 | 0.140 | 0.711 | 0.01 |
| | | T25 | 1 | 1 | 23.30 | 1.341 | 0.259 | 0.05 | 1 | 1 | 23.61 | 0.376 | 0.546 | 0.02 |
| | Beta 1 | T0 | 1 | 1 | 22.84 | 0.287 | 0.597 | 0.01 | 1 | 1 | 21.88 | 0.002 | 0.966 | 0.00 |
| | | T15 | 1 | 1 | 24.66 | 0.020 | 0.888 | 0.00 | 1 | 1 | 24.85 | 0.174 | 0.680 | 0.01 |
| | | T25 | 1 | 1 | 24.28 | 0.586 | 0.451 | 0.02 | 1 | 1 | 24.34 | 0.586 | 0.451 | 0.02 |
| Sham | Alpha | T0 | 1 | 1 | 19.57 | 0.155 | 0.698 | 0.01 | 1 | 1 | 21.55 | 0.316 | 0.580 | 0.01 |
| | | T15 | 1 | 1 | 21.53 | 0.032 | 0.859 | 0.00 | 1 | 1 | 23.08 | 0.003 | 0.961 | 0.00 |
| | | T25 | 1 | 1 | 23.41 | 0.527 | 0.475 | 0.02 | 1 | 1 | 24.14 | 0.779 | 0.386 | 0.03 |
| | Beta 1 | T0 | 1 | 1 | 24.13 | 0.414 | 0.526 | 0.02 | 1 | 1 | 24.49 | 0.000 | 0.989 | 0.00 |
| | | T15 | 1 | 1 | 24.34 | 0.006 | 0.938 | 0.00 | 1 | 1 | 24.38 | 0.218 | 0.645 | 0.01 |
| | | T25 | 1 | 1 | 24.31 | 0.019 | 0.892 | 0.00 | 1 | 1 | 23.91 | 0.015 | 0.903 | 0.00 |

DF – degrees of freedom for the numerator, DF Den – degrees of freedom for the denominator, p-value (uncorr) – uncorrected p-value, ROI Left and ROI Right – region of interest in the left and right hemisphere

Supplementary table S5. Results of linear mixed model analysis of TBS-induced modulation of absolute EEG power in ROI Left and ROI Right across visits

| Protocol | Frequency | Time postTBS | ROI Left | | | | | | ROI Right | | | | | |
|---|---|---|---|---|---|---|---|---|---|---|---|---|---|---|
| | | | N parm | DF | DF Den | F-value | p-value (uncorr) | Partial eta | N parm | DF | DF Den | F-value | p-value (uncorr) | Partial eta |
| cTBS | Alpha | T0 | 1 | 1 | 22.93 | 0.549 | 0.466 | 0.02 | 1 | 1 | 23.32 | 0.361 | 0.554 | 0.02 |
| | | T15 | 1 | 1 | 21.90 | 0.114 | 0.738 | 0.01 | 1 | 1 | 22.29 | 0.015 | 0.905 | 0.00 |
| | | T25 | 1 | 1 | 23.39 | 0.003 | 0.960 | 0.00 | 1 | 1 | 23.36 | 0.004 | 0.950 | 0.00 |
| | Beta 1 | T0 | 1 | 1 | 22.16 | 0.000 | 0.987 | 0.00 | 1 | 1 | 22.88 | 0.538 | 0.471 | 0.02 |
| | | T15 | 1 | 1 | 22.94 | 1.299 | 0.266 | 0.05 | 1 | 1 | 22.61 | 1.435 | 0.243 | 0.06 |
| | | T25 | 1 | 1 | 23.18 | 0.700 | 0.412 | 0.03 | 1 | 1 | 22.97 | 1.222 | 0.281 | 0.05 |
| iTBS | Alpha | T0 | 1 | 1 | 22.78 | 0.386 | 0.541 | 0.02 | 1 | 1 | 22.73 | 0.094 | 0.763 | 0.00 |
| | | T15 | 1 | 1 | 23.37 | 0.000 | 0.989 | 0.00 | 1 | 1 | 23.33 | 0.120 | 0.732 | 0.01 |
| | | T25 | 1 | 1 | 24.15 | 0.021 | 0.887 | 0.00 | 1 | 1 | 24.03 | 0.217 | 0.646 | 0.01 |
| | Beta 1 | T0 | 1 | 1 | 21.96 | 0.313 | 0.581 | 0.01 | 1 | 1 | 22.60 | 0.789 | 0.384 | 0.03 |
| | | T15 | 1 | 1 | 20.27 | 0.061 | 0.808 | 0.00 | 1 | 1 | 17.71 | 0.427 | 0.522 | 0.02 |
| | | T25 | 1 | 1 | 22.96 | 0.351 | 0.560 | 0.02 | 1 | 1 | 23.76 | 0.595 | 0.448 | 0.02 |
| Sham | Alpha | T0 | 1 | 1 | 22.67 | 0.796 | 0.382 | 0.03 | 1 | 1 | 23.24 | 0.058 | 0.811 | 0.00 |
| | | T15 | 1 | 1 | 23.25 | 0.237 | 0.631 | 0.01 | 1 | 1 | 23.70 | 0.020 | 0.890 | 0.00 |
| | | T25 | 1 | 1 | 24.32 | 0.083 | 0.775 | 0.00 | 1 | 1 | 24.05 | 0.117 | 0.735 | 0.00 |
| | Beta 1 | T0 | 1 | 1 | 20.71 | 0.694 | 0.415 | 0.03 | 1 | 1 | 19.28 | 0.426 | 0.522 | 0.02 |
| | | T15 | 1 | 1 | 21.56 | 1.578 | 0.223 | 0.07 | 1 | 1 | 21.35 | 0.486 | 0.493 | 0.02 |
| | | T25 | 1 | 1 | 22.91 | 0.062 | 0.805 | 0.00 | 1 | 1 | 22.66 | 0.006 | 0.942 | 0.00 |

DF – degrees of freedom for the numerator, DF Den – degrees of freedom for the denominator, p-value (uncorr) – uncorrected p-value, ROI Left and ROI Right – region of interest in the left and right hemisphere

Supplementary Table S6. Cronbach's alpha coefficients of consistency of TBS-induced EEG modulation across visits

| Protocol | Frequency band | Time postTBS | Relative power | Absolute power | |
|---|---|---|---|---|---|
| | | | ROI Right | ROI Left | ROI Right |
| cTBS | Alpha | T0 | 0.504 | 0.045 | 0.028 |
| | | T15 | 0.677 | 0.744 | 0.691 |
| | | T25 | 0.668 | 0.216 | 0.304 |
| | Beta 1 | T0 | 0.174 | 0.021 | 0.303 |
| | | T15 | 0.829 | 0.262 | 0.588 |
| | | T25 | 0.378 | 0.296 | 0.498 |
| iTBS | Alpha | T0 | 0.067 | 0.286 | 0.276 |
| | | T15 | 0.748 | 0.475 | 0.499 |
| | | T25 | 0.310 | 0.443 | 0.436 |
| | Beta 1 | T0 | 0.332 | 0.127 | 0.068 |
| | | T15 | 0.531 | 0.365 | 0.509 |
| | | T25 | 0.494 | 0.338 | 0.392 |
| Sham | Alpha | T0 | 0.027 | 0.262 | 0.346 |
| | | T15 | 0.006 | 0.067 | 0.493 |
| | | T25 | 0.712 | 0.691 | 0.852 |
| | Beta 1 | T0 | 0.684 | 0.460 | 0.601 |
| | | T15 | 0.777 | 0.674 | 0.390 |
| | | T25 | 0.980 | 0.346 | 0.258 |

Supplementary Table S7. Results of contingency analysis of individual TBS-induced changes

| Protocol | Frequency | Relative power | | | | | Absolute power | | | | |
|---|---|---|---|---|---|---|---|---|---|---|---|
| | | N | DF | Pearson chi-square | p-value (uncorr.) | Type conversion (%) | N | DF | Pearson chi-square | p-value (uncorr.) | Type conversion (%) |
| cTBS | Alpha | 45 | 3 | 3.989 | 0.263 | 76 | 45 | 3 | 3.966 | 0.265 | 57 |
| | Beta 1 | 45 | 3 | 1.752 | 0.417 | 67 | 45 | 3 | 2.312 | 0.510 | 71 |
| iTBS | Alpha | 43 | 3 | 3.354 | 0.340 | 79 | 43 | 3 | 1.556 | 0.669 | 79 |
| | Beta 1 | 43 | 3 | 2.244 | 0.523 | 63 | 43 | 3 | 6.226 | 0.101 | 79 |
| Sham | Alpha | 44 | 3 | 0.470 | 0.925 | 65 | 44 | 3 | 1.079 | 0.782 | 65 |
| | Beta 1 | 44 | 3 | 0.369 | 0.832 | 55 | 44 | 3 | 4.978 | 0.173 | 40 |

N – number of samples, DF – degrees of freedom, p-value (uncorr.) – uncorrected p-value, type conversion – number of participants demonstrating different types of EEG change across two visits.

Supplementary table S8. Numerical proportions of types of TBS-induced changes of absolute EEG power in ROI Left in the initial (V1) and retest (V2) visits

| Protocol | Visit | Alpha | | | | Beta 1 | | | |
|---|---|---|---|---|---|---|---|---|---|
| | | Increase of power | Decrease of power | Mixed change | No change | Increase of power | Decrease of power | Mixed change | No change |
| cTBS | V1 | 48 | 30 | 9 | 13 | 35 | 35 | 9 | 22 |
| | V2 | 27 | 55 | 14 | 5 | 36 | 45 | 0 | 18 |
| iTBS | V1 | 24 | 43 | 10 | 24 | 19 | 24 | 10 | 48 |
| | V2 | 36 | 45 | 5 | 14 | 36 | 41 | 9 | 14 |
| Sham | V1 | 29 | 33 | 10 | 29 | 33 | 10 | 10 | 48 |
| | V2 | 43 | 26 | 9 | 22 | 43 | 26 | 0 | 30 |

Supplementary table S9. Results of linear mixed model analysis of TBS-induced modulation of relative EEG power in ROI Left and ROI Right across active and sham TBS protocols

| Visit | Frequency | Factor | ROI Left | | | | | | ROI Right | | | | | |
|---|---|---|---|---|---|---|---|---|---|---|---|---|---|---|
| | | | N parm | DF | DF Den | F-value | p-value (uncorr) | Partial eta | N parm | DF | DF Den | F-value | p-value (uncorr) | Partial eta |
| V1 | Alpha | Protocol | 2 | 2 | 166.6 | 2.018 | 0.136 | 0.02 | 2 | 2 | 165.1 | 1.595 | 0.206 | 0.02 |
| | | Protocol x Time | 4 | 4 | 157.0 | 0.815 | 0.518 | 0.02 | 4 | 4 | 155.6 | 0.417 | 0.796 | 0.01 |
| | Beta 1 | Protocol | 2 | 2 | 167.3 | 2.104 | 0.125 | 0.02 | 2 | 2 | 170.2 | 4.685 | **0.011** | 0.05 |
| | | Protocol x Time | 4 | 4 | 158.8 | 0.353 | 0.841 | 0.01 | 4 | 4 | 159.9 | 0.302 | 0.876 | 0.01 |
| V2 | Alpha | Protocol | 2 | 2 | 173.9 | 0.783 | 0.459 | 0.01 | 2 | 2 | 175.1 | 0.077 | 0.926 | 0.00 |
| | | Protocol x Time | 4 | 4 | 169.3 | 0.149 | 0.963 | 0.00 | 4 | 4 | 170.1 | 0.140 | 0.967 | 0.00 |
| | Beta 1 | Protocol | 2 | 2 | 174.9 | 1.462 | 0.235 | 0.02 | 2 | 2 | 176.1 | 1.414 | 0.246 | 0.02 |
| | | Protocol x Time | 4 | 4 | 170.1 | 0.404 | 0.806 | 0.01 | 4 | 4 | 170.6 | 0.149 | 0.963 | 0.00 |

DF – degrees of freedom for the numerator, DF Den – degrees of freedom for the denominator, p-value (uncorr) – uncorrected p-value, ROI Left and ROI Right – region of interest in the left and right hemisphere

Supplementary table S10. Results of linear mixed model analysis of TBS-induced modulation of absolute EEG power in ROI Left and ROI Right across active and sham TBS protocols

| Visit | Frequency | Factor | ROI Left | | | | | | ROI Right | | | | | |
|---|---|---|---|---|---|---|---|---|---|---|---|---|---|---|
| | | | N parm | DF | DF Den | F-value | p-value (uncorr) | Partial eta | N parm | DF | DF Den | F-value | p-value (uncorr) | Partial eta |
| V1 | Alpha | Protocol | 2 | 2 | 168.5 | 1.060 | 0.349 | 0.01 | 2 | 2 | 167.6 | 1.834 | 0.163 | 0.02 |
| | | Protocol x Time | 4 | 4 | 158.5 | 0.714 | 0.584 | 0.02 | 4 | 4 | 158 | 0.623 | 0.647 | 0.02 |
| | Beta 1 | Protocol | 2 | 2 | 165.2 | 0.808 | 0.448 | 0.01 | 2 | 2 | 164.3 | 1.293 | 0.277 | 0.02 |
| | | Protocol x Time | 4 | 4 | 158.4 | 0.612 | 0.655 | 0.02 | 4 | 4 | 158.3 | 0.967 | 0.428 | 0.02 |
| V2 | Alpha | Protocol | 2 | 2 | 174.6 | 2.948 | 0.055 | 0.03 | 2 | 2 | 175.8 | 1.617 | 0.201 | 0.02 |
| | | Protocol x Time | 4 | 4 | 170.4 | 0.311 | 0.870 | 0.01 | 4 | 4 | 170.7 | 0.191 | 0.943 | 0.00 |
| | Beta 1 | Protocol | 2 | 2 | 173.3 | 4.813 | **0.009** | 0.05 | 2 | 2 | 173.6 | 3.308 | **0.039** | 0.04 |
| | | Protocol x Time | 4 | 4 | 170.1 | 0.768 | 0.547 | 0.02 | 4 | 4 | 169.9 | 0.358 | 0.839 | 0.01 |

DF – degrees of freedom for the numerator, DF Den – degrees of freedom for the denominator, p-value (uncorr) – uncorrected p-value, ROI Left and ROI Right – region of interest in the left and right hemisphere
